# A proposed microcosm for landscape ecology – beyond the binary to the patch-mosaic model

**DOI:** 10.1101/542985

**Authors:** Yolanda F. Wiersma, Rachel D. Wigle, R. Troy McMullin

## Abstract

**Background:** Microcosms such as pitcher plants, or patches of mosses on a rock surface, have been used worldwide to allow for manipulative experiments that test hypotheses for patterns observed at larger extents, such as dispersal or community assemblage. Such microcosms can also be applied to questions in landscape ecology, but are limited by their binary (patch/non-patch structure). Here we examine a more realistic model landscape system that shares the patch-mosaic structure common to kilometres-extent landscapes. This system of lichen thalli on tree trunks has been shown to have consistent spatial patterns across replicate microcosms, but only when sampling within a limited area. To be relevant for experimentation across scales, it is necessary to determine whether previously observed patterns are consistent when sampling across a broader region and when using different tree species. Here, we test for consistent landscape patch pattern in both maco- and micro-lichens across 21 balsam fir (*Abies balsamea*) and yellow birch (*Betula alleghaniensis*) trees.

**Methods:** We measured spatial pattern of lichen thalli along the trunks of two species of trees, at two spatial resolutions; trees within a single stand (∼100 × 50 m) and trees dispersed across a larger region (500 km^2^). We used a “lichen ladder” comprised of 5 10 × 10 cm sampling blocks to quantify number of species and individuals in a 50 cm section of the tree trunk. We tested for similar patterns along the trunk and between the north and south sides at both sampling intensities using perMANOVA.

**Results:** We find that lichen patches on tree trunks can function as replicate microscoms for landscape ecology. Patterns of thalli along the trunks of trees and between the north and south aspects of the trunk are statistically significantly consistent, although there is variation between tree species, and groups of lichens included. Our microcosm could be used as a model system for landscape ecology research; but researchers should test for consistent patterns first.

## Introduction

Research in ecology has always been challenged with the trade-off between experimental rigour and real-world applicability (Schindler 1998; Kohler 2002). The tenets of good experimental design (control, randomization and replication) are done most easily through manipulative experiments in a lab setting, while observational studies allow for natural processes to remain present, at the expense of the experimental rigour (Kohler 2002). A widely-used solution to bridge this divide is through the use of mesocosms (Stewart et al. 2013) or microcosms (Srivastava et al. 2004). *Mesocosms* and *microcosms* (see Box 1 for glossary) share the property of being contained systems, usually small enough to allow for adequate replication for statistical power and that can be experimentally manipulated. The distinction between the two is that mescocosms are artificially constructed while microscoms (the focus of this paper) occur naturally (Box 1). Commonly used microcosms include pitcher plants (e.g., Mouquet et al. 2008), bromeliads (e.g., Talaga et al. 2015), patches of mosses (e.g., Gonzalez et al. 1998) and rocky tidepools (e.g., Schiesari et al. 2018). These are sometimes termed “*microecosystems*” in both lab (Drake 1993) and field (Gilbert et al. 1998; Lau et al. 2018) contexts; however in this paper we will take natural microecoystems as equivalent to microcosms (Box 1). In a review, Srivastava et al. (2004) suggest that natural microcosms are potentially highly powerful tools that can advance ecological research questions but caution that the utility of microcosms is limited to the types of questions applied (Srivastava et al. 2004). In addition, Srivastava et al. (2004) and others (Zuk and Travisano 2018) emphasize that microcosms should be viewed as complementary tools to theoretical models, lab experiments and field observations.

**Box 1. Glossary of Terms**

*mesocosm*: a contained system (e.g., aquarium) that is designed to mimic a natural, larger ecosystem for the purposes of experimental manipulation.

*microcosm*: a small, contained natural ecosystem, such as insects in a pitcher plant, or a tide pool community.

*microecosystem*: a small ecosystem that is constrained/bounded by specific structural or environmental factors (e.g., pH, substrate).

*patch*: the fundamental unit in a landscape; a patch is considered to be homogenous within, relative to the landscape and have its own intrinsic properties (e.g., size, shape).

*landscape*: an areas that is spatially heterogeneous in at least one factor of interest.

*landscape mosaic*: a landscape comprised of two or more patches (ecosystems) that interact to create a pattern at a larger extent.

*hierarchy*: a conceptual framework for natural systems; lower levels of organization (finer grain/smaller extent) are constrained by higher levels, yet at the same time, lower levels influence higher level patterns. For example, gaps are nested within forest stands; stands are nested within tracts and tracts within a larger ecosystem which is nested within a biome.

Ecological research using microcosms has mainly focused on questions in community and population ecology and has successfully used them as a tool to examine topics such as dispersal (e.g., Grainger and Gilbert 2016), predator-prey dynamics (e.g., Busse et al. 2019) and community assemblage (e.g., Pereira et al. 2018). Although the microcosm as a whole is often referred to as a “landscape” (Drake 1993; Gilbert et al. 1998), the experimental systems described above are all simple binary landscapes of habitat and non-habitat patches (i.e., pitcher plant leaves and the surrounding air, tide pools and the surrounding rocks). Even though experimental manipulation of moss patches (Gilbert et al. 1998; Gonzalez et al. 1998; Gonzalez and Chaneton 2002) contained three species of bryophytes, these different species were treated as functionally equivalent patches in the experiments. Such simplification can be appropriate, depending on the research question; however a binary patch/non-patch landscape is an over-simplification of the way landscape ecologists view the world (Box 1), and may explain why microcosms have mainly been used for questions in population and community ecology and not yet expanded widely to landscape ecology (but see Jenerette and Shen 2012).

Landscape ecologists commonly view their object of study using the “patch-mosaic” model, that is, that *landscapes* are comprised of a diversity of *patch* types, each of which has its own intrinsic properties of size, shape and cover, but which interacts with adjacent patterns to form a *landscape mosaic* (Box 1; Wu 2013). Thus, while the binary microcosms of moss patches, bromeliads or tide pools have been shown to be good model systems for some processes that occur at larger extents (e.g., species dispersal and assembly on oceanic islands as predicted by island biogeography theory), they may not function well as model systems for more complex landscapes.

There have not been many developments of microcosms for landscape ecology experiments, although experimental mesocosms have been proposed as a tool (e.g., Wiens and Milne 1989). Model systems that have been proposed for landscape ecology include soil biocrusts (Bowker et al. 2014) and lichen-covered tree trunks (Wiersma and McMullin 2018). Bowker et al. (2014) point out that soil biocrusts are most like scaled-down landscapes in that they have similar attributes to kilometers-extent landscapes, such as patch heterogeneity. Moreover, their small size makes them tractable for manipulative experiments with good potential for experimental replicates, and they can also be collected and transferred into controlled environments (e.g., greenhouses) for increased experimental rigour. At the same time, they are interesting ecological systems independent of their use as model systems, which is an often-overlooked property of models (Zuk and Travisano 2018). Wiersma and McMullin (2018) proposed a similar system of heterogeneous patches of cryptogams as Bowker et al. (2014), but growing along the boles of trees as replicate landscapes. Although Bowker et al.’s (2014) system and Wiersma and McMullin’s (2018) share similar types of organisms and patch pattern, Wiersma and McMullin’s (2018) has the additional feature of being explicitly hierarchical (Fig. 1) with the disadvantage relative to Bowker et al.’s (2014) of not being transferrable to a lab setting.

**Figure 1.**
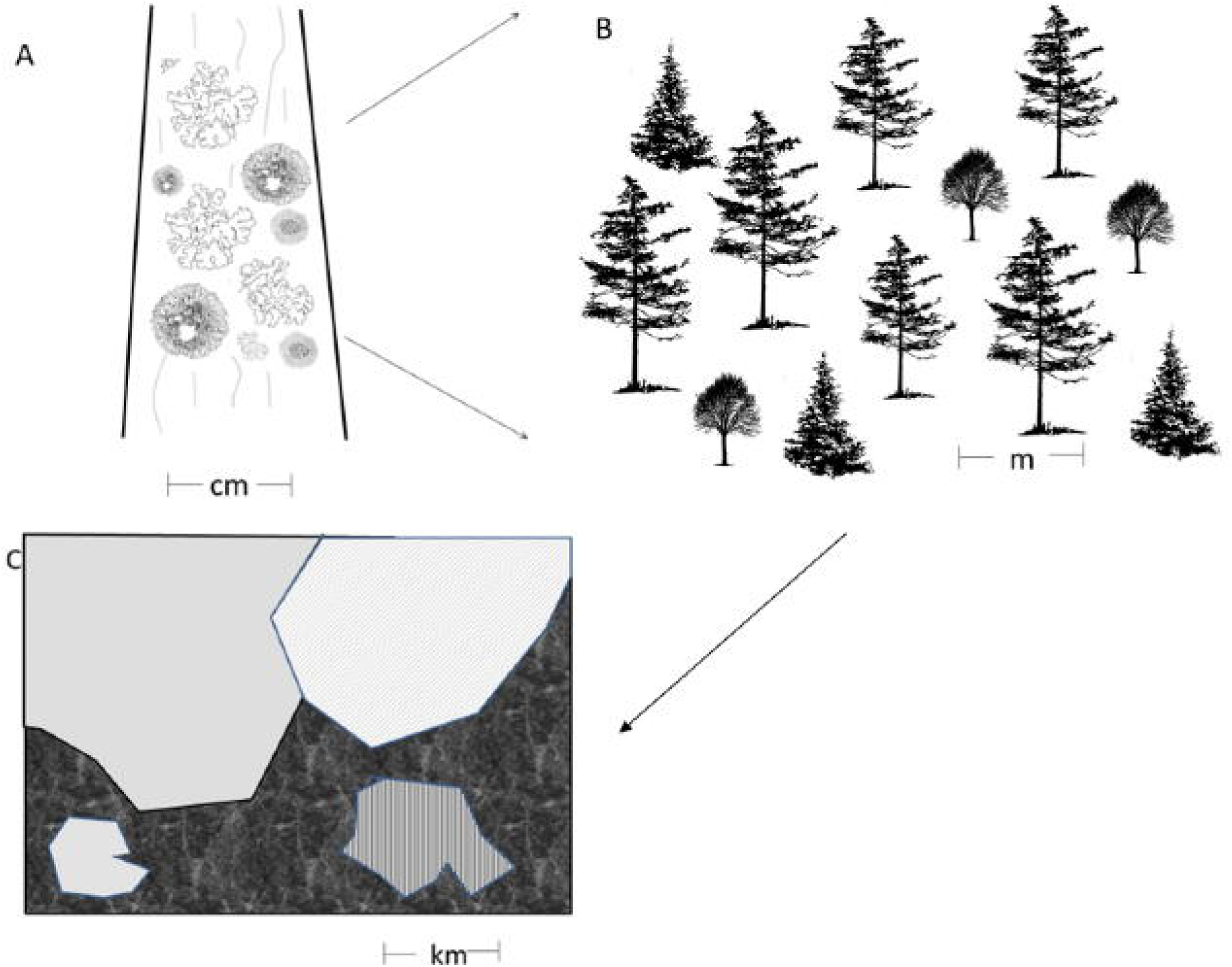
A schematic of the hierarchical structure of the proposed model system for landscapes. At the smallest scale (panel a) the landscape is centimeters in extent and is comprised of patches of lichens on the bole for trees. Each tree can be considered a replicate microlandscape. Within a single forest stand (panel b), the landscape is metres in extent. Some stands are homogenous in terms of tree species, age/size, but across the stand there will be fine-grained heterogeneity in microtopography and microclimate. These stands are then patches in a kilometers-extent landscape (panel c); some stands of similar composition may be replicated across the landscape but interspersed with other patch types (e.g., non-forested bog, meadow, anthropogenic patches).

*Hierarchy* is a key concept in landscape ecology (Urban et al. 1987; O’Neill et al. 1989; Kotliar and Wiens 1990) and it has been suggested that by harnessing principles of hierarchy and scaling (Urban et al. 1987; Urban 2005; Bowker et al. 2014), that microlandscape patterns and processes (a level at which experimentation is more feasible) can be extrapolated to larger extents (a level at which many management challenges occur). Wiersma and McMullin (2018) demonstrated that 2 sets of 12 balsam fir (*Abies balsamea*) trees within a very small area (< 100 m apart) share similar patch patterns of macro-lichens along the boles, and proposed that these trees could be considered replicate (micro)landscapes. While this system holds promise as a potential new type of microcosm for landscape ecology, the limited focus of a single species of tree in a restricted geographic area limits the broader applicability of their proposed model system. Here, we use additional data – from two species of trees and across a much wider extent (500 km^2^) and with a broader suite of lichens, to scrutinize Wiersma and McMullin’s (2018) hypothesis that patches of lichens along a tree bole are a model system for landscapes. If the patch-patterns along the boles of trees holds true in this expanded study, then we posit that similar systems in other regions might also function as complex landscape microcosms that could be harnessed for experimentation.

## Methods

### Study area

We carried out sampling in the Avalon Forest Ecoregion on the island of Newfoundland, Canada. The Avalon Forest is the smallest (500 km^2^) ecoregion in the province of Newfoundland and Labrador, characterized by high humidity and precipitation, cool summers, and mild winters (South 1983). The forested areas are on rolling hills called ribbed moraines, landscape features created by glaciers (Hättestrand and Kleman 1999). Interspersed between the moraines are open, sphagnum-dominated wetlands (South 1983). Forests are dominated by balsam fir (*Abies balsamea*) with black spruce (*Picea mariana*) in wet areas and occasional yellow birch (*Betula alleghaniensis*) stands on north facing slopes (South 1983).

Field work was carried out under Permit Nos. 2015/16-14 and 2017/18-10 issued by the Department of Environment and Conservation, Government of Newfoundland and Labrador.

### Experimental Design

We visited 22 sites across the region (Fig. 2) and at 21 of these, selected two trees, one balsam fir and one yellow birch that were similar in diameter and within 25 m of each other. The remaining site (indicated as “Halls Gullies” on Fig. 2) was the original location intensively sampled (24 trees total) by Wiersma and McMullin (2018). All sampling locations were balsam-fir dominated stands, mostly occurring on moraines. On each tree, we sampled on the north and south sides of the bole using a 10 cm × 50 cm “lichen ladder”, divided into five 10 × 10 cm “blocks”, placed from 1.1 m to 1.6 m up the bole (Fig. 3). We identified and counted all lichen species within each 10 cm block. Species that could not be identified in the field were collected for identification using standard processes, including microscopy, chemical spot tests (Brodo et al. 2001), and thin-layer chromatography (Culberson and Kristinsson 1970). We inventoried both macro-(those with a more three-dimensional growth from, and growing on the substrate, which includes foliose and fruticose growth forms) and micro-lichens (those growing within the substrate, i.e., crustose growth forms) at the 21 new sites; Wiersma and McMullin (2018) limited their study to field-identifiable macro-lichens. Thus, we analyzed the data from the 21 dispersed sites with macro-lichens only to compare to the previous study; we also repeated the analysis for more dispersed trees using data on both macro- and micro-lichens to assess whether the landscape patterns observed by Wiersma and McMullin (2018) held when examining a broader suite of lichen species and when looking across a larger sampling area.

**Figure 2.**
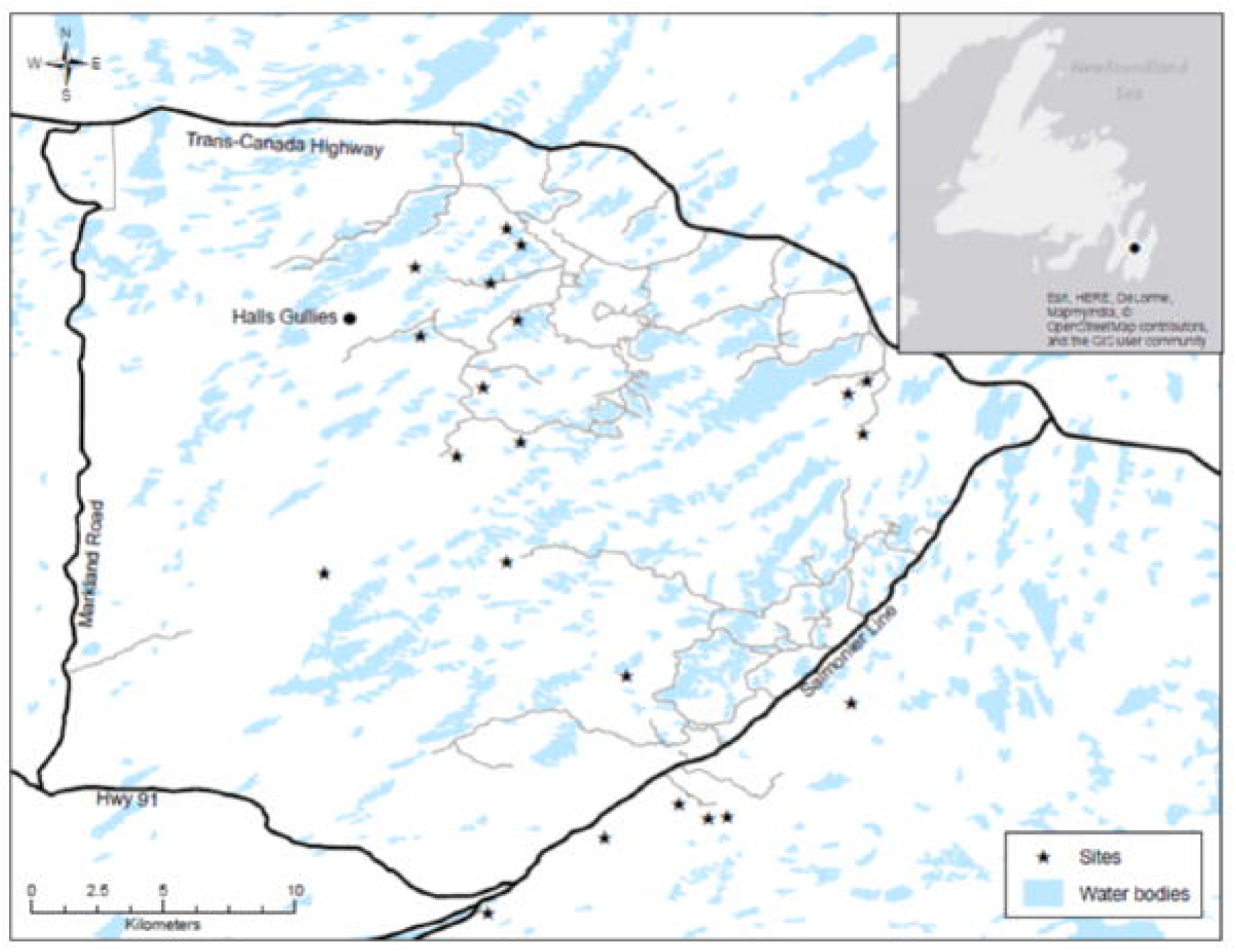
Location of sampling sites in the Avalon Forest Ecoregion on the island of Newfoundland, Canada. Inset map shows the location of the Avalon Forest Ecoregion (black polygon). Black stars on the main map are sampling locations for the 21 sites where we sampled both balsam fir and yellow birch. The location labelled “Halls Gullies” designates the stand within which 24 balsam fir were sampled.

**Figure 3.**
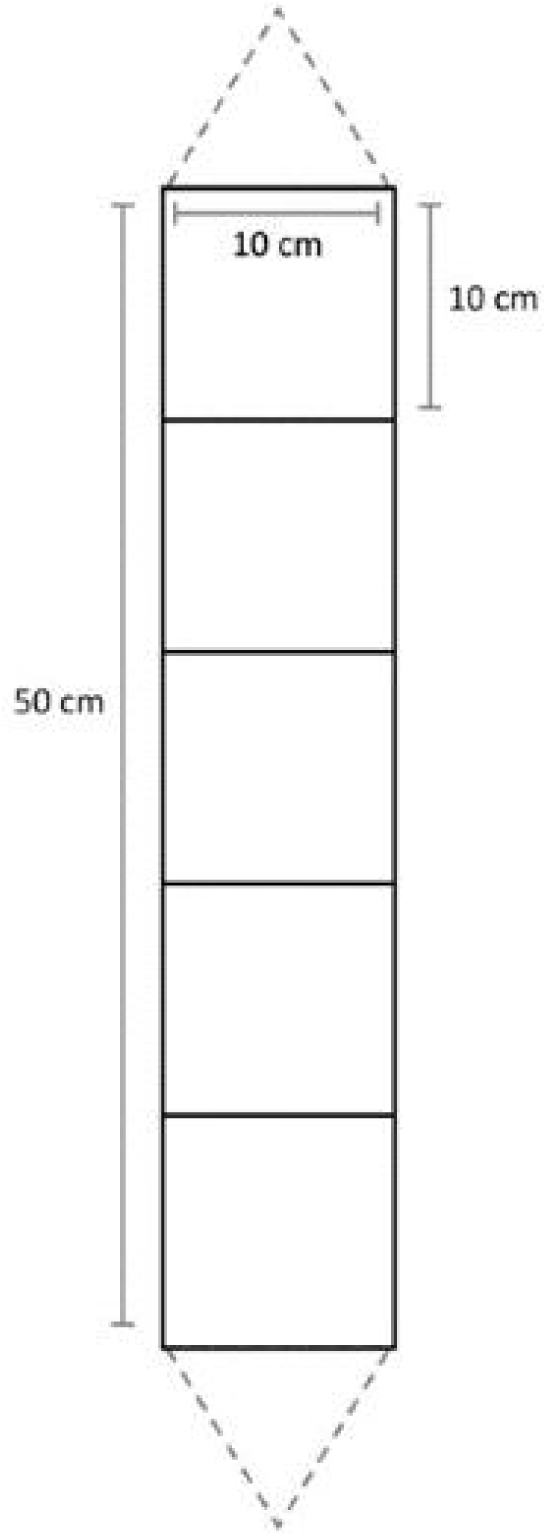
A lichen ladder, which was used to sample lichen diversity on tree boles. Dimensions of each square are 10 × 10 cm; the ladder is 50 cm in length and was placed with the top rung at 1.6 m from the ground.

### Statistical analysis

We used a perMANOVA analysis (Anderson 2001) to assess whether the pattern of lichen patches along the bole was consistent across all trees when stratifying for aspect, and whether the patterns between the north and south sides were consistent when stratifying by height up the bole. We separately analysed the data from the 21 balsam fir and the 21 yellow birch trees in the expanded study area and did not combine the data from the 21 balsam fir in the wider ecoregion with the data from the 24 balsam fir in the single stand because of differences in geographic sampling intensity. We carried out all statistical analysis using R (version 1.0.136; R Core Team 2016) with the package vegan (Oksanen et al. 2016).

## 3. Results

The 24 balsam fir trees in the single stand showed a significantly consistent lichen patch pattern between the north and south sides along the 50 cm portion of the bole, when stratifying by tree (perMANOVA R^2^ = 0.00552, *p* = 0.05) but not when we controlled for position along the bole (perMANOVA R^2^ = 0.00552, *p* = 0.223). There was also a significant pattern for position along the bole when controlling for the tree (perMANOVA R^2^ = 0.001727, *p* = 0.024) but not for position along the bole when we stratified for aspect (perMANOVA R^2^ = 0.01727, *p* = 0.412).

The perMANOVA results for the 21 more spatially dispersed balsam fir trees also showed a significant pattern for macro-lichens between the north and south sides of the tree, when stratifying by tree (perMANOVA R^2^ = 0.00988, *p* = 0.009) but not when controlling for position along the bole (perMANOVA R^2^ = 0.0098, *p* = 0.081). Unlike for the trees in the single stand, there was no significant pattern for position along the bole when controlling for the tree (perMANOVA R^2^ = 0.00476, *p* = 0.157) nor for position along the bole when stratified for aspect (perMANOVA R^2^ = 0.00476, p = 0.444). When examining a different tree species, yellow birch, there was no significantly consistent pattern of lichen between the north and south sides of the bole, either when stratifying by tree (perMANOVA R^2^ = 0.00147, *p* = 0.772) or by position along the bole (perMANOVA R^2^ = 0.00147, *p* = 0.85). Nor was there any significant pattern along the bole of the yellow birch when stratifying by tree (perMANOVA R^2^ = 0.00143, *p* = 0.721) or when controlling for aspect (perMANOVA R^2^ = 0.00143, *p* = 0.811). Overall patterns for macro-lichens are summarized in Table 1.

**Table 1.**
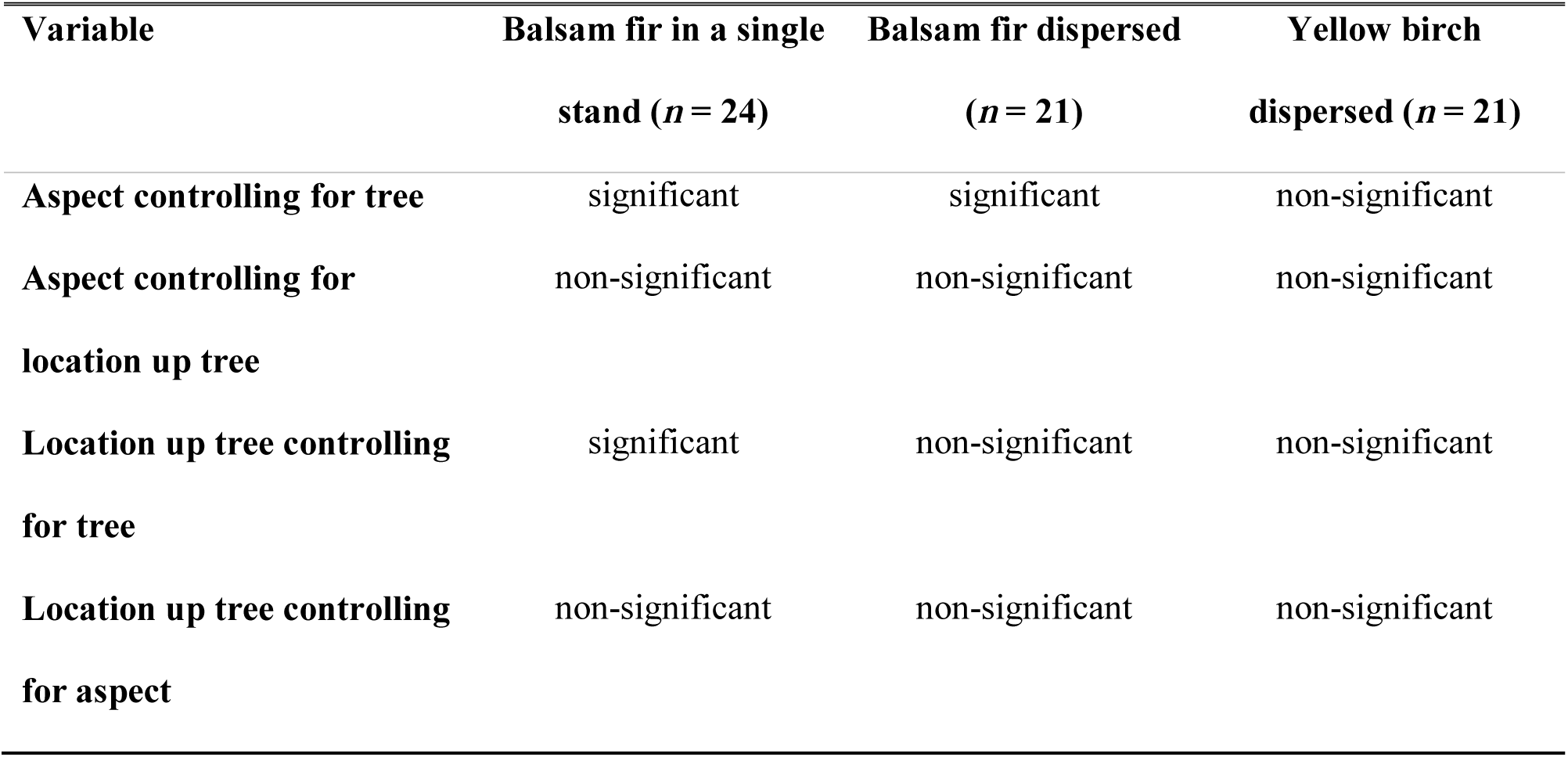
Summary of perMANOVA analysis for trees within the Avalon Forest Ecoregion. Date is from two sampling schema; one for 24 balsam fir in a single stand; and for 21 more widely dispersed sites with one balsam fir and one yellow birch at each site. The analysis is for consistency in lichen patterns on the tree boles <u>for macro-lichens only</u> along a 50 cm section of the bole on the north- and south-facing sides.

| Variable | Balsam fir in a single<br>stand (*n* = 24) | Balsam fir dispersed<br>(*n* = 21) | Yellow birch<br>dispersed (*n* = 21) |
| --- | --- | --- | --- |
| Aspect controlling for tree | significant | significant | non-significant |
| Aspect controlling for<br>location up tree | non-significant | non-significant | non-significant |
| Location up tree controlling<br>for tree | significant | non-significant | non-significant |
| Location up tree controlling<br>for aspect | non-significant | non-significant | non-significant |

When we looked at both macro- and micro-lichens, the patterns were different. Balsam fir did not show any significant pattern (aspect stratified by tree perMANOVA R^2^ = 0.00497 *p* = 0.051; aspect stratified by position along bole perMANOVA R^2^ = 0.00497, *p* = 0.439; position along bole stratified by tree perMANOVA R^2^ = 0.00245, *p* = 0.471; position along bole stratified by aspect perMANOVA R^2^ = 0.00245, *p* = 0.843). In contrast, there was a significant pattern for aspect for yellow birch, both when stratifying by tree (perMANOVA R^2^ = 0.01528, *p* = 0.001) and by position along the bole (perMANOVA R^2^ = 0.01528, *p* = 0.002). However, there was not a significant pattern for location up the bole for yellow birch, neither when stratifying by tree (perMANOVA R^2^ = 0.00373, *p* = 0.297) nor by aspect (perMANOVA R^2^ = 0.00373, *p* = 0.647). Table 2 summarizes the overall patterns for macro- and micro-lichens combined.

**Table 2.**
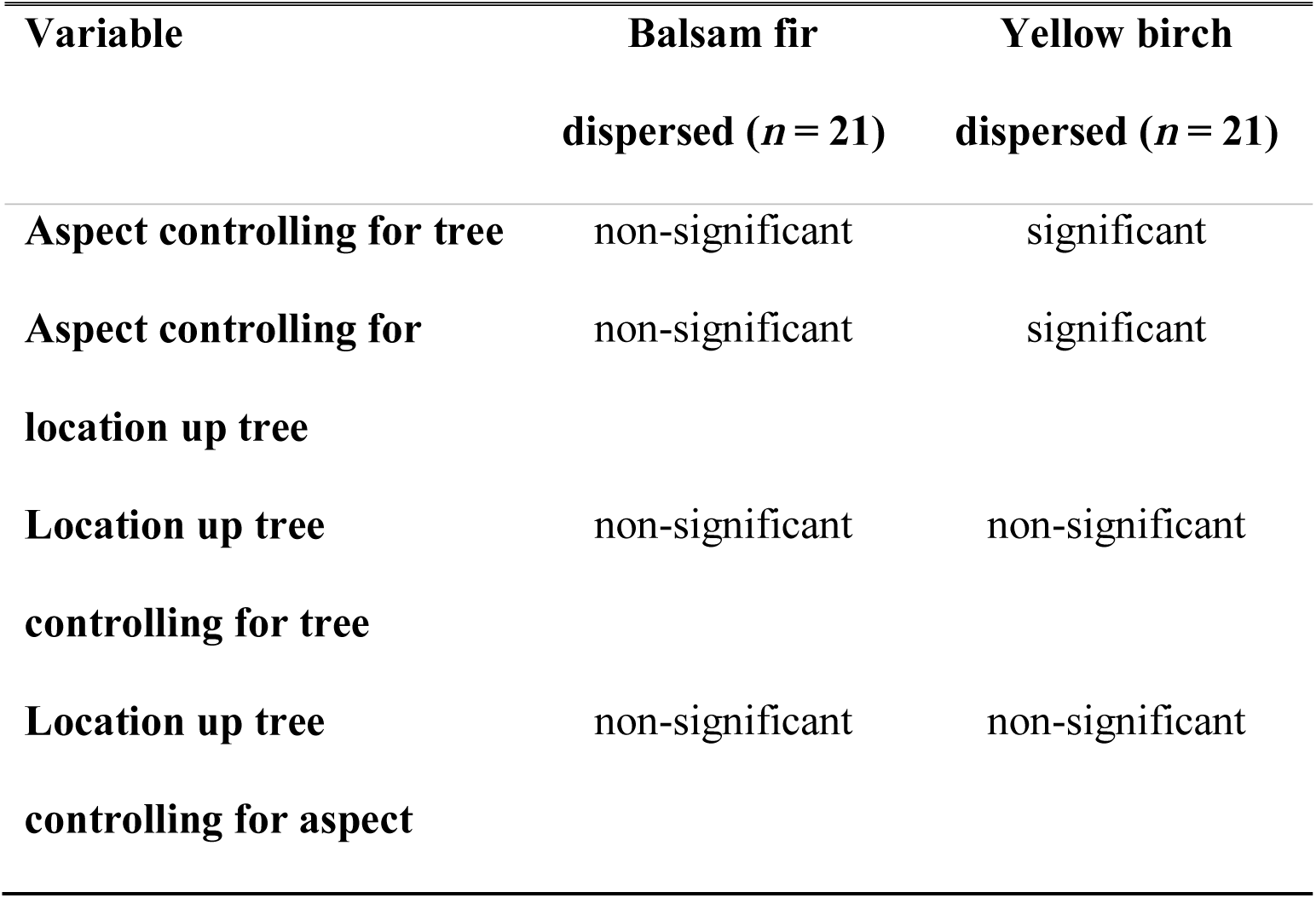
Summary of perMANOVA analysis for 21 widely dispersed sites across the Avalon Forest Ecoregion. Data are shown for balsam fir and yellow birch; there was one of each at each site. The analysis is for consistency in lichen patterns along 50 cm of the tree boles along the north and south-facing sides, <u>for macro-lichens and mirco-lichens combined</u>.

## Discussion

The consistent patterns of patches of lichens growing along tree boles, even when trees are widely dispersed (and presumably exposed to different micro-climate and stand-level constraints) suggest that these microecosystems can be considered a model system for landscape ecology. Macro-lichen patch pattern on the north vs. south sides of the bole of balsam fir trees that were widely dispersed followed that of the more aggregated trees. Although the pattern along the gradient of the bole was non-significant for the dispersed trees, we believe this is due to the fact that the microtransect we used here was only 50 cm in length. Wiersma and McMullin (2018) detected consistent patch patterns along a 1 metre gradient of a tree bole, and (Young et al. 2018) documented gradients in both epiphyte diversity and microfauna along gradients running 25 m up Douglas fir (*Pseudotsuga menziesii*) trees. Landscape patterns and gradients in kilometer-extent landscapes are only detected when measured at an appropriate scale, and likely 50 cm along the bole was insufficient to robustly detect any gradient in the 21 dispersed trees. We did, however, detect a statistically significantly consistent pattern along a 50 cm gradient of the bole in the 24 more clumped trees (where Wiersma and McMullin (2018) detected an even stronger pattern along a 1 m gradient), which increases our confidence that a pattern does exist in the more dispersed trees, we were simply unable to detect it.

While macro-lichens growing on the balsam fir trees appear to have consistent patch patterns, and thus suggest these as microcosms amenable to experimentation, this proposed system should be approached cautiously. Macro-lichens along the bole of yellow birch trees did not show any consistent pattern. However, when we included both macro- and micro-lichens, yellow birch did show consistent patch patterns on the north vs. south sides of the boles. Conversely, patterns on balsam fir disappeared when both macro- and micro-lichens were included. Thus, researchers wishing to adopt systems of cryptogams growing along tree boles should first assess which community (i.e., just macro-lichens, or macro- and micro-lichens combined), growing on which tree species, has a consistent spatial pattern and hence can function as a landscape microcosm.

Despite that not all lichens growing on both species of trees showed a consistent spatial pattern, we believe this microcosm of lichens along a tree bole holds promise for research. As with other natural microcosms, this system allows for experimental replicates to test different kinds of ecological questions. The complexity of multiple patch types and patch patterns allows for different kinds of questions than those afforded by the binary patch/non-patch landscapes of most microcosm work to date. Some questions that could be addressed include the spatial distribution of microfauna (e.g., Bokhorst et al. 2015; Trekels et al. 2017; Young et al. 2018) or more complex “island biogeography” questions (e.g., Southwood and Kennedy 1983). Srivastava et al. (2004) suggested four future research questions that could be addressed with natural microcosms; the patch patterns of lichen-covered tree trunks would be especially relevant to question about how species characteristics and interactions drive patterns at spatial scales (Srivastava et al. 2004). The hierarchical nature of this proposed model system will also yield insights into how ecological processes change with system size (Srivastava et al. 2004). Research on meta-communities is an exciting and expanded area (Logue et al. 2011), but often lacks a spatially explicit approach. The addition of a microcosm with a spatially explicit pattern adds a novel tool to both the community and landscape ecologist’s tool box.

Potential limitations to this system are that, unlike the soil crusts proposed as a model system for community and landscape ecology by Bowker et al. (2014), trees cannot be transported to a lab for experimentation under controlled conditions. Moreover, the slow growth of lichens may make the response time to certain manipulations (e.g., experimental fragmentation) too slow to be practical for most studies. However, some lichens are relatively fast growing (Phillips 1969; Faegri 1980; Bidussi and Gauslaa 2015) and thus by carefully choosing manipulations, this system has the potential to contribute a novel type of microcosm that replicates the patterns and processes observed (but not amenable to experimentation) in kilometers-extent landscapes.

Our finding here, that spatial patterns in a natural microcosm are reasonably consistent for trees that are widely dispersed, and that the landscape patch-mosaic model appears to be hierarchical, has advantages for other kinds of research questions. Previously, Wiersma and McMullin (2018) exploited these microcosms in a single, spatially-constrained stand to test whether particular features of these micro-landscapes influenced presence or absence of a rare species. In this case, controlling for stand-level constraints was helpful to eliminate other larger-extent factors that might influence rare species persistence. However, the ability to replicate microcosms across a broader region allows for natural-experiments to see how landscapes respond to meso-scale conditions such as distance to different types of habitat (e.g., open bogs), or broader gradients in elevation and climate. It also allows for manipulative experiments at larger extents that might mimic real-world processes, for example, looking at responses to different disturbance levels.

## Conclusion

Thinking of the microcosm of epiphytes growing along a tree trunk as a model system for landscapes opens up a whole suite of research opportunities not available in the binary microcosms that have been the norm to date. The replicated patch-pattern allows for these microcosms to be treated as micro-landscapes and thus have replicate landscape units to test spatially-explicit questions; replication that is not possible with kilometers-extent landscapes. Such a system is not amenable, or even necessary, for all types of research questions; parsiomony should dictate when a binary landscape is sufficient for hypothesis testing. At the same time, our system is interesting in and of itself; something that Zuk and Travisano (2018) emphasize should not be overlooked with model systems. Micro-fauna and flora are often incredibly diverse, and not always well-understood. Thus, we encourage further research on these fascinating microcosms which occur in forests worldwide.

## Acknowledgements

Thanks are due to T. Sato, P. Lauriault, J. Godfrey, M. Fahmy, E. Kissler, T. Padgett, G. Wachinger, M.Wilkes and C. Ziter for assistance with field work, and to E. Lopez and M. Fahmy for data entry. Salmonier Nature Park graciously supplied housing to RTM for the 2015 field work.

